# Genome-wide RNA structural determination in *Candida* yeast pathogens

**DOI:** 10.1101/2023.12.27.573417

**Authors:** Uciel Chorostecki, Ester Saus, Toni Gabaldón

## Abstract

Understanding the intricate roles of RNA molecules in virulence and host-pathogen interactions can provide valuable insights into combatting infections and improving human health. Although much progress has been achieved in understanding transcriptional regulation during host-pathogen interactions in diverse species, more is needed to know about the structure of pathogen RNAs. This is particularly true for fungal pathogens, including pathogenic yeasts of the *Candida* genus, which are the leading cause of hospital-acquired fungal infections. Deciphering the relation between RNA structure and their biology remains a significant gap. Despite advancements in transcriptional regulation studies, especially for fungal pathogens like *Candida*, the structural aspects of pathogenic RNAs remain understudied. Our work addresses this gap by employing genome-wide structure probing to comprehensively explore the structural landscape of mRNAs and long non-coding RNAs (lncRNAs) in the four major *Candida* pathogens. Specifically focusing on mRNA, we observe a robust correlation between sequence conservation and structural characteristics in orthologous transcripts, significantly when sequence identity exceeds 50%, highlighting structural feature conservation among closely related species. We investigate the impact of single nucleotide polymorphisms (SNPs) on mRNA secondary structure. SNPs within 5’ untranslated regions (UTRs) tend to occur in less structured positions, suggesting structural constraints influencing transcript regulation. Furthermore, we compare the structural properties of coding regions and UTRs, noting that coding regions are generally more structured than UTRs, consistent with similar trends in other species.

Additionally, we provide the first experimental characterization of lncRNA structures in *Candida species*. Most lncRNAs form independent subdomains, similar to human lncRNAs. Notably, we identify hairpin-like structures in lncRNAs, a feature known to be functionally significant. Comparing hairpin prevalence between lncRNAs and protein-coding genes, we find enrichment in lncRNAs across *Candida* species, humans, and *Arabidopsis thaliana*, suggesting a conserved role for these structures.

In summary, our study offers valuable insights into the interplay between RNA sequence, structure, and function in *Candida* pathogens, with implications for gene expression regulation and potential therapeutic strategies against *Candida* infections.

## Introduction

Yeasts belonging to the *Candida* genus are among the most common human fungal opportunistic pathogens (Brown et al. 2012). Up to 30 distinct, phylogenetically diverse *Candida* species have been reported to infect humans, and the risk of infection is especially high in immunocompromised hospitalized patients (Gabaldón, Naranjo-Ortíz, and Marcet-Houben 2016; Papon et al. 2013). Although their incidence may differ from region to region, the four most common species, *Candida albicans*, *Candida glabrata*, *Candida parapsilosis* and *Candida tropicalis*, generally in this order, collectively account for over 95% of the cases *(Diekema et al. 2012),* all of them included in the fungal priority pathogens list, by the world health organization (World Health Organization. Antimicrobial Resistance Division et al. 2022).

The transcriptome comprises all the RNA transcribed by the genome in a specific cell type or tissue, at a particular developmental stage, and under a specific condition (Jacquier 2009). Thus, transcriptome analysis allows for understanding the genome’s activity and provides comprehension of gene expression regulation, structure and function. Accumulating data overwhelmingly supports the idea that any RNA’s function is always determined by its structure (Vicens and Kieft 2022). For instance, recent studies have shown that the regulation of mRNA stability through RNA modification is a crucial step for achieving a tight regulation of gene expression (Delaunay and Frye 2019), and mRNA stability depends on the mRNA nucleotide sequence, which affects the secondary and tertiary structures of the mRNAs. Several methods were developed for the computational prediction of RNA secondary structure, and the performance of the methods varies across the datasets (Bugnon et al. 2022).

Moreover, the accuracy of existing algorithms still needs to be improved to model long RNA molecules since the number of possible structures scales dramatically with the length of the sequence (Wan et al. 2012; Bugnon et al. 2022). The structure prediction accuracy is improved either by searching for conservation in a set of homologous sequences or by using experimental data. In this regard, several experimental methods have been developed to study RNA structure at single-nucleotide resolution (Saus et al. 2018; Kertesz et al. 2010; Wan et al. 2013; Underwood et al. 2010).

The structural content of the UTRs compared to the coding regions has been found to vary from organism to organism (Mortimer, Kidwell, and Doudna 2014). Secondary structures of 5’ UTR, the coding region, and 3’ UTR are mainly independent since base pairs across domain boundaries are infrequent (Shabalina, Ogurtsov, and Spiridonov 2006). 5ʹ UTR structures in eukaryotic mRNAs can modulate the translation initiation. Moreover, regulatory elements in the mRNA, especially in the 3ʹ UTR, can also modulate translation (Leppek, Das, and Barna 2018). It is important to note that even within 5’ UTRs, unstructured (linear) regulatory elements are likely to have a crucial impact on translation (Hinnebusch, Ivanov, and Sonenberg 2016). The median length of mRNA 5ʹ UTRs in humans is around 200 nt, exceeding those of other mammals and tripling that of budding yeast (about 50 nt) (Leppek, Das, and Barna 2018).

Sequence-structure relationships in mRNA coding regions remain elusive, and their secondary structure is largely unknown. The correlation between sequence and structure in *Saccharomyces cerevisiae* mRNAs coding regions is weaker than in sncRNAs, and profiles of paralogous mRNAs show a strong correlation with sequence for identity levels upwards of 85–90% (Chursov et al. 2011). However, pairs of more distantly related yeast transcripts’ secondary structures appear to be unrelated (Chursov et al. 2011). Furthermore, in a previous study, the structures of orthologous mRNAs from *Saccharomyces cerevisiae* and *C. glabrata* were studied only *in-silico*. That study found no correlation for pairs with low sequence identity levels, and there was no conclusion for similar sequences due to a lack of data (Chursov et al. 2011).

The human body temperature is a robust defence against fungal infections, particularly in cases of fever (Bergman and Casadevall 2010). High temperatures significantly limit fungal growth. During Candida infections, the host’s fever response exposes fungal cells to temperatures between 37°C and 42°C. These temperature changes profoundly impact various aspects of *Candida*, such as its appearance, reproduction, characteristics, and drug resistance (Shapiro, Robbins, and Cowen 2011). This insight can guide strategies for combating *Candida* infections. Moreover, RNA secondary structures exhibit high sensitivity to temperature variations and modifying temperature conditions can induce alterations in RNA folding and stability. Exploring RNA structures across various temperatures unveils dynamic structural changes that may play a pivotal role in governing gene expression, especially in response to temperature shifts encountered during infection processes.

Long non-coding RNAs (lncRNAs) represent a heterogeneous group of ncRNAs, longer than 200 nucleotides (Ponting, Oliver, and Reik 2009), and it remains unclear the function of most of them. These molecules exhibit distinct characteristics consistently observed across diverse taxa, including mammals, insects, and plants. Compared to protein-coding genes, they tend to have lower expression levels and greater cell type specificity (Cabili et al. 2011). Their sequence conservation is generally poor, and they undergo rapid evolutionary changes, frequently displaying species-specific traits (Kutter et al. 2012). Recent research sheds light on the significance of lncRNAs in *Candida* species. In a comprehensive study encompassing five major *Candida* pathogens, hundreds of lncRNAs were identified from publicly available sequencing data (Hovhannisyan and Gabaldón 2021). These lncRNAs exhibit unique evolutionary characteristics, and despite limited sequence conservation among these species, some lncRNAs share common sequence motifs and show co-expression with specific protein-coding transcripts, suggesting potential functional connections.

Understanding RNA structure is vital as it can help uncover the regulation of mRNAs and the roles of lncRNAs and even reveal potential functional consequences of synonymous or non-coding genetic variations. Thus, to gain further insights into the role of RNA structure in *Candida* species, we performed RNA probing experiments and comparative genomics analysis of five yeast species, including the four major *Candida* species and the model yeast *S. cerevisiae*. To our knowledge, our study constitutes the most comprehensive examination of structures of mRNAs and lncRNAs in yeasts. We determined and compared pairs of orthologs across the considered species to shed new light on the relationships between sequence and structure conservation. In addition, for one of the species, we compared the RNA structures at varying temperatures and measured their relative stabilities. Finally, we reached the nextPARS score across the coding sequence (CDS) and untranslated regions (UTRs) in *C. glabrata*. Our results show that the coding regions in *C. glabrata* transcripts are more structured than untranslated regions.

Our study provides a comprehensive analysis of mRNA and lncRNA structures in the major *Candida* pathogens, shedding light on the conservation of structural features and their correlation with sequence conservation. Candida species have developed resistance mechanisms to antifungal agents, making treatment increasingly challenging. Understanding the intricate relationship between sequence, structure, and function will contribute to understanding the evolution of *Candida* species, potentially identifying new strategies to combat *Candida* infections.

## Methods

### Sample preparation, secondary structure probing with NextPARS and sequencing

We performed four different experiments to probe the secondary structure of transcriptomes of four *Candida* species: *C. albicans* SC5314, *C. parapsilosis* GA1, *C. glabrata* CBS138 and *C. tropicalis* CSPO. Cultures were set up for each species and were grown in YPD medium in an orbital shaker at 200 rpm, overnight, at 37°C for *C.glabrata* and 30°C for the rest of the species. Total RNA was extracted from these cultures using the RiboPure™-Yeast Kit according to the manufacturer’s instructions (ThermoFisher Scientific), starting with a total amount of 3x10^8^ cells per sample as recommended for a maximum yield. To obtain PolyA+ RNA samples, total RNA from yeast were purified by two rounds of selection using Dynabeads mRNA purification kit following the manufacturer’s instructions (ThermoFisher Scientific). The quality (RNA integrity) and quantity of both total and PolyA+ RNA samples were assessed using the Agilent 2100 Bioanalyzer with the RNA 6000 Nano LabChip Kit (Agilent), the NanoDrop 1000 Spectrophotometer (ThermoFisher Scientific), and the Qubit Fluorometer with the Qubit RNA BR (Broad-Range) Assay Kit (ThermoFisher Scientific).

We used the NextPARs approach (PMID: 29358234, PMID: 34141143), to probe the secondary structure of the transcriptomes at 23 °C, 37 °C and 55 °C for *C. parapsilosis* and at 23°C for the remaining species. Two μg of the corresponding polyA+ RNA were used per each reaction tube. 0.03 U of RNase V1 (Ambion) and 200 U of S1 nuclease (Fermentas) were used to digest the corresponding samples when probing the structure at 23°C. When samples were probed at 37°C and 55°C, the final enzyme concentration was adapted to prevent overdigestion of RNAs due to higher temperatures, according to previous studies (PMID: 22981864). Thus, in the digestion reactions, 0.015 U or 0.0075 U of Rnase V1 (Ambion), and 100 U or 50 U of S1 nuclease (Fermentas) were used to probe the transcriptomes at 37°C or 55°C, respectively. We performed quality controls of the final digested samples to confirm that they were not over-digested after enzymatic treatment.

Once the good quality of the final digested samples was confirmed, libraries were prepared using the TruSeq Small RNA Sample Preparation Kit (Illumina) following a modified protocol previously described (Saus et al. 2018; Chorostecki, Saus, and Gabaldón 2021). After performing quality control of each library using Agilent 2100 Bioanalyzer with the DNA 1000 kit (Agilent), libraries were sequenced in single-reads with read lengths of 50 nucleotides in Illumina HiSeq2500 sequencers at the Genomics Unit of the CRG (CRG-CNAG).

For *S. cerevisiae*, we used available data previously generated by the nextPARS technique (Bioproject id PRJNA380612) (Saus et al. 2018).

### Secondary structure prediction

First, we converted the nextPARS score to SHAPE-like reactivities with the nextPARS2SHAPE v1.0 script (https://github.com/Gabaldonlab/MutiFolds/blob/master/scripts/nextPARS2SHAPE.py), and we used it for the secondary structure prediction. Then, to obtain the secondary structure of *Candida* mRNAs and lncRNAs, we use the Fold software (Version 6.2) from the RNAstructure package (Reuter and Mathews 2010), using pseudo energy restraints. Residues for which no nextPARS data were assigned a reactivity of 999, as suggested by the Fold manual. Additionally, we calculated the similarity score for lncRNAs folded with and without nextPARS data using the R package RNAsmc (Wang et al. 2023). This package provides a similarity score for two RNA secondary structures, with larger values indicating greater similarity between the structures. The score ranges from 0, representing no similarity, to a maximum value of 10, signifying identical RNA secondary structures.

### Stem-loop prediction

We retrieve the sequence information of the four *Candida* species, human and *Arabidopsis thaliana.* Then, to obtain the secondary structure of protein-coding genes (PCG) and lncRNAs, we use the Fold software (Version 6.2) from the RNAstructure package (Reuter and Mathews 2010). Finally, the stem-loop prediction was made by developing an in-house script (stem-loop_predictor_public.py) using forgi (version 2.0.0), a Python library for manipulating RNA secondary structure. The folding and the search for stem-loop in different species have been performed on MareNostrum 4 supercomputer at the Barcelona Supercomputing Center, Spain. The code is available on GitHub (https://github.com/Gabaldonlab/Candida_nextPARS).

### RNA secondary structure prediction and visualization

To obtain the secondary structure of the lncRNA and PCG, we use RNAstructure software (Reuter and Mathews 2010). Using the nextPARS2SHAPE.py script (https://github.com/Gabaldonlab/MutiFolds/), we converted the nextPARS score to SHAPE-like normalized reactivities that were used to provide pseudoenergy restraints to the Fold software. For residues for which there was no nextPARS data, we assigned a reactivity of 999, as suggested by the RNAfold manual. The RNA arc diagram was built using R-chie (version 2.0.) (Tsybulskyi, Mounir, and Meyer 2020). RNA structures were constructed using VARNA (Version 3-93) (Darty, Denise, and Ponty 2009).

### Shuffled analysis for sequence-structure correlation

We performed a shuffled analysis as a control to elucidate further the relationship between sequence conservation and structural features in orthologous transcripts across yeast species. For that, we randomly shuffled the positions of nextPARS scores and their alignments for orthologous pairs of genes retrieved from different species. We then calculated the correlation of the nextPARS scores for these shuffled positions in the same way for the real data. This approach allowed us to assess whether the observed sequence-structure correlations in orthologous transcripts were significantly higher than those expected by random chance.

### Statistical methods

To compare nextPARS data among *C. parapsilosis* mRNAs at various temperatures, the non-parametric Kruskal-Wallis test was employed, considering the non-normal distribution of the data. Similarly, the assessment of nextPARS scores between UTRs and coding regions in *C. glabrata* involved the application of the Kruskal-Wallis test, accommodating the non-normality of the dataset. Furthermore, in the analysis focusing on SNP positions within the 5’ UTR in *C. glabrata*, a paired T-test was utilized for the comparison, considering the paired nature of the data.

### SNP analysis

To investigate the impact of secondary structure induced by SNPs in *Candida* transcripts, we used variant calling data obtained from CandidaMine (https://candidamine.org/), an integrative data warehouse for *Candida*. We focused on identifying SNPs within the coding regions of genes for all four *Candida* species studied. We compared the nextPARS scores in loci with identified SNPs to those in loci with no reported SNPs. Additionally, for *C. glabrata*, we extended our analysis to the untranslated regions (UTRs), as these regions remain unannotated in other *Candida* species. Specifically, we assessed how SNPs within the 5’ UTRs of *C. glabrata* transcripts correlate with RNA secondary structure, focusing on identifying any potential structural changes associated with these SNPs. We developed custom Python (version 3.10.9) scripts (compare_score_SNP.py) using the pandas library (version 2.1.1) to process and compare the SNPs and nextPARS scores. The code is available on GitHub (https://github.com/Gabaldonlab/Candida_nextPARS). Specifically, we assessed how SNPs within the 5’ UTRs of *C. glabrata* transcripts correlate with RNA secondary structure, focusing on identifying any potential structural changes associated with these SNPs. Here, a threshold of 0.2 was applied to enhance precision and minimize noise. This threshold was explicitly implemented to reduce potential interference from positions with a score of 0, which could arise due to the absence of nextPARS data at those particular positions.

### Calculation of Long-Range Interactions

The percentage of long-range interactions in the secondary structure of each lncRNA sequence was determined using a custom Python script. The script parses the dot-bracket notation, representing RNA secondary structure, and identifies base pairs. A dynamically calculated threshold based on a specified percentage of the sequence length is employed to define long-range interactions. We used a default threshold of 25%, indicating that interactions with a base pair distance exceeding 25% of the sequence length are considered long-range. Hence, we compute the percentage of long-range interactions and generate a plot illustrating the variation across different lncRNAs.

## Data availability

The raw sequencing data of nextPARS experiments used in this project have been deposited in the Short Read Archive of the European Nucleotide Archive under the Bioproject IDs PRJNA714002 and PRJNA838569.

## Results

### Genome-wide view of the structural landscape of Candida mRNAs

To compare the structural landscape of *Candida* RNAs, we performed nextPARS experiments (Saus et al. 2018; Chorostecki et al. 2021) in the four most commonly infecting *Candida* species: *C, albicans*, *C. glabrata*, *C. parapsilosis,* and *C. tropicalis*, which collectively account for over 95% of all Candida infections (Consortium OPATHY and Gabaldón 2019). NextPARS yields direct measures of *in vitro* secondary structure at single-nucleotide resolution. This information is represented by a structural profile providing a score for each residue that ranges from -1 (highest preference for single strand) to 1 (highest preference for double strand). After filtering out transcripts with low counts we obtained structural information on mRNAs and lncRNAs (see Materials and Methods, Supplementary Table 1). This dataset, comprising experimental probing information from hundreds of transcripts, provides the first genome-wide view of the structural landscape of *Candida* mRNAs and lncRNAs. For comparative purposes, we also used, in downstream analyses, publicly available nextPARS data for the model yeast *Saccharomyces cerevisiae (Saus et al. 2018)*.

### Assessing sequence-structure relationships in Candida mRNAs

There needs to be a better understanding of how sequence conservation relates to the conservation of structural features in orthologous transcripts across different species. We measured the sequence and structure similarity correlation in mRNAs coding regions to assess this for yeast species. We retrieved pairs of orthologous genes between the five species using the MetaPhOrs (v2.5) web server (Chorostecki et al. 2020). For all pairs of orthologous mRNAs with nextPARS information (Supplementary Figure 1), we calculated the correlation of the nextPARS score in aligned positions. We took different sequence identity intervals and observed high correlations of nextPARS scores (median Pearson correlation > 0.4) when the sequence percent identity is above ∼50% (Figure 1A). This value is significantly higher than expected by chance, as correlations between randomly shuffled scores were consistently below 0.2 (Supplementary Figure 2). Levels of structural conservation across orthologous pairs reflected phylogeny (Gabaldón, Naranjo-Ortíz, and Marcet-Houben 2016), where CTG *Candida* species (*C. albicans, C. parapsilosis, C. tropicalis*) and post-WGD species (*C. glabrata* and *S. cerevisiae*) belong to two distant clades (Figure 1B).

**Figure 1.**
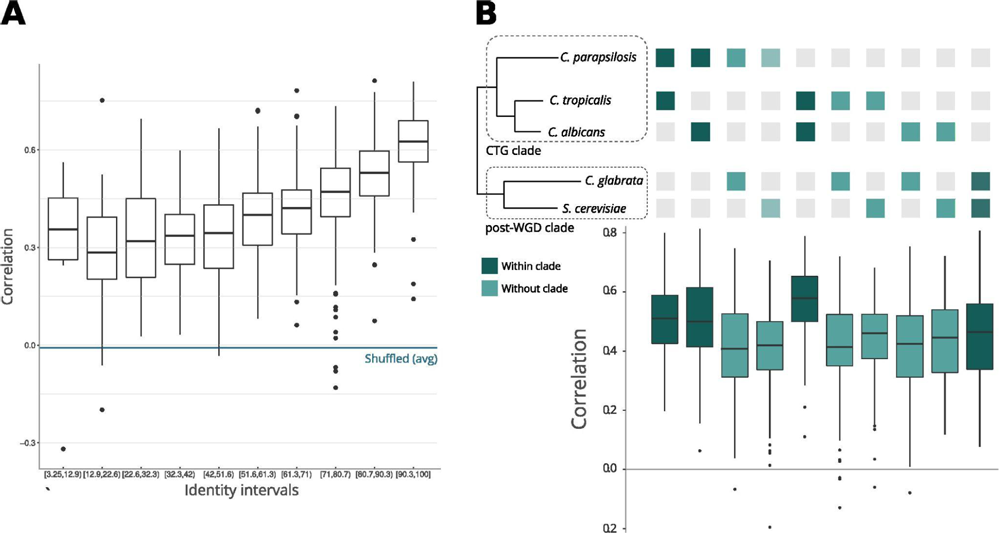
Comparison of sequences and structures across yeast species. **(A)** Pair sequence alignments by intervals. The X-axis represents sequence identity intervals as a percentage, while the Y-axis illustrates the correlations of nextPARS scores across various identity intervals. **(B)** Correlation between sequence and structure in orthologs. The orthologs represent pairs of genes from the species depicted in the tree below the graph. The tree’s topology and evolutionary distances are adapted from (Gabaldón, Naranjo-Ortíz, and Marcet-Houben 2016).

Moreover, we compared the correlations of the nextPARS score in aligned positions but using multiple sequence alignments from orthologs in the five species, using each species as a seed (Supplementary Figure 1). On average, we observed a positive correlation between the nextPARS score in aligned positions (Supplementary Figure 3).

Investigating alterations in transcript secondary structure induced by temperature changes is crucial, as it helps elucidate how *Candida* pathogens dynamically regulate their gene expression in response to temperature shifts encountered during infection processes. Thus, to investigate the alterations in transcript secondary structure prompted by temperature changes in *C. parapsilosis* mRNAs, we performed nextPARS experiments at three different temperatures: 23 °C, 37 °C and 55 °C. We noticed differences on average in the nextPARS score at different temperatures (Supplementary Figure 4A), where the significant differences are between 23 °C compared with 37 °C and 55 °C. We also measured the temperature stability at different temperatures but found no significant difference when comparing conserved positions against non-conserved ones (Supplementary Figure 4B).

### Exploring structural dynamics in C. glabrata mRNA regions in Coding Sequences vs UTRs and the Impact of SNPs

Exploring the structural characteristics of coding and untranslated regions in *C. glabrata* mRNAs is pivotal for understanding the intricacies of gene expression regulation. Thus, we investigate the structure in different regions within mRNAs. We compared the average nextPARS score across the coding sequence (CDS) and untranslated regions (UTRs) in *C. glabrata*. CDS exhibit significantly higher nextPARS scores than 5’ and 3’ UTRs (Figure 2A; p-value < 1.7 e-06 and p-value < 0.00053, respectively). These findings agree with the results observed in other species, such as *Saccharomyces cerevisiae* and *Arabidopsis thaliana (Kertesz et al. 2010; Li, Zheng, Vandivier, et al. 2012)*. However, opposite results have been observed in humans, *Drosophila melanogaster* and *Caenorhabditis elegans,* in which CDS are slightly more single-stranded than the UTRs (Wan et al. 2014; Li, Zheng, Ryvkin, et al. 2012).

**Figure 2.**
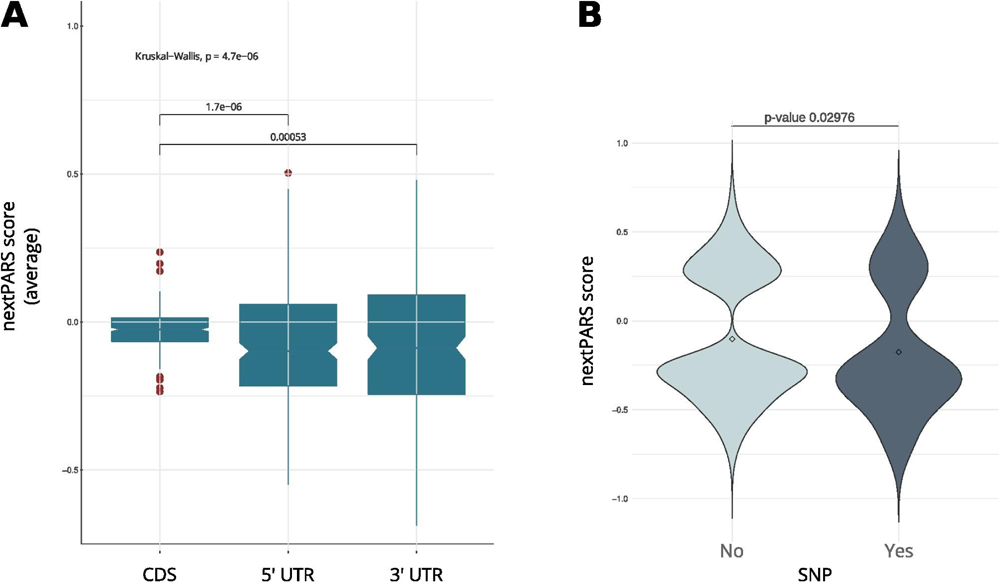
RNA structure comparative analysis in *C. glabrata.* **(A)** A boxplot showing a comparison between coding regions in *C. glabrata* and UTRs. **(B)** The nextPARS score in SNP vs. no SNP positions in *C glabrata ’*s 5’ UTR.

We then delved into the impact of secondary structure on sequence variation, analyzing SNPs in UTRs from *C. glabrata*. We used variant calling data obtained from CandidaMine (https://candidamine.org/) - an integrative data warehouse for *Candida* yeasts. We compared nextPARS scores in loci with SNPs to those in loci with no reported SNPs. Our results show no differences between those two groups for any of the four *Candida* species (Supplementary Figure 5). Then, we performed a similar analysis using UTRs from *C. glabrata* - as these features remain unannotated in the other *Candida species*. Changes in those regions are associated with deregulation in gene expression at transcriptional and post-transcriptional levels (Lawless et al. 2009). When we compared SNP’s positions in UTRs, we observed that SNPs in 5’ UTR tended to localize to less structured positions (Figure 2B; p-value < 0.02976), shedding light on potential regulatory regulation in these critical regions. However, in the 3’ UTR, a difference in means was noted, albeit with limited statistical significance (p-value = 0.2491), due to the small number of values in SNP positions. This observation underscores the dynamic interplay between sequence variations and structural features, adding depth to our understanding of the intricate regulatory landscape within *Candida* yeasts.

### Structural characterization of Candida lncRNAs using experimental information

Catalogues of long non-coding RNAs (lncRNAs) in major *Candida* species have only been recently characterized (Hovhannisyan and Gabaldón 2021). We used our nextPARS data to characterize the secondary structure of inferred lncRNAs in the four *Candida* species considered in this study. Given the generally low expression levels of lncRNAs, we could only detect a few lncRNAs with sufficient confidence (> five average counts per position) (Supplementary Table 1). These nevertheless provide the first structural insights of lncRNAS in these relevant pathogens. Furthermore, it is crucial to highlight that our characterization of lncRNAs involved a unique approach. Leveraging the experimental insights provided by our nextPARS data, we employed these data as constraints during the RNA folding process of lncRNAs. This approach allowed us to guide *in-silico* secondary structure predictions with the experimental data. We noticed significant differences for certain lncRNAs compared to predictions made without these constraints (see Materials and Methods, Supplementary Figure 6). Moreover, we observed that most lncRNAs folded in potential independent subdomains with few long-range interactions (see Materials and Methods, Supplementary Figure 7) in accordance with previous studies on human lncRNAs (Somarowthu et al. 2015; Ziv et al. 2021; Chorostecki, Saus, and Gabaldón 2021).

Next, we focused on a specific lncRNA known as MSTRG.4167.1, which displayed upregulation in C. albicans during epithelial cell infection, as reported by Hovhannisyan et al. (Hovhannisyan and Gabaldón 2021). To investigate its structural characteristics, we folded the lncRNA using data from nextPARS experiments as constraints (Figure 3A). We examined this particular lncRNA in detail based on the experimental data we obtained through our nextPARS experiments. We compared predicted RNA structures using data from nextPARS as constraints for experiments conducted at different temperatures for this particular lncRNA. By utilizing similarity scores of the structures, we observed that significant differences emerged when comparing MSTRG.4167.1 at 23 and 37 degrees, with an even more substantial distinction between 37 and 55 degrees. The most pronounced difference was when comparing the lncRNA at 23 and 55 degrees (Supplementary Table 2). As expected, the similarity score decreases as the temperature difference increases. These results imply that the lncRNA folds differently at distinct temperatures, which could result in varied functions or regulations. Furthermore, we noticed a slight trend of fewer long-range interactions for MSTRG.4167.1 as the temperature increased (Supplementary Figure 8).

**Figure 3.**
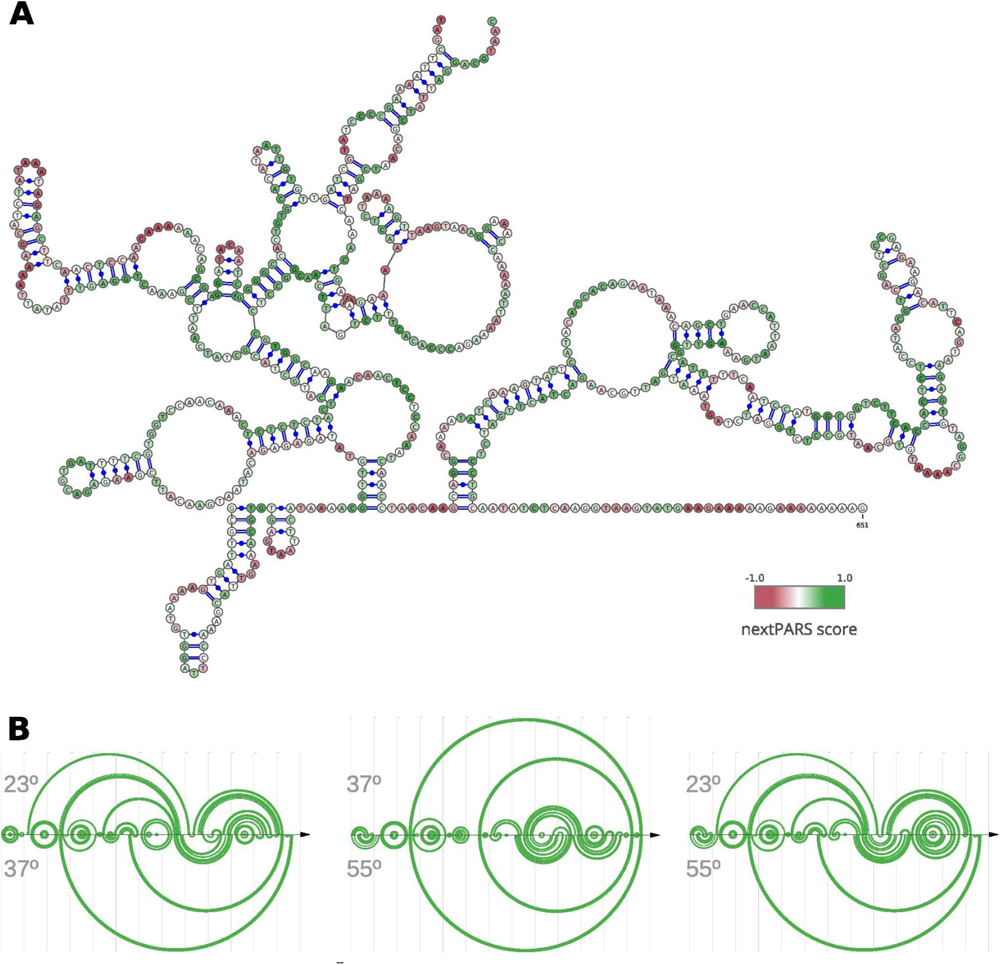
Structural Dynamics of lncRNA MSTRG.4167.1. This figure illustrates the structure of a specific lncRNA, MSTRG.4167.1. **(A)** RNA secondary structure of MSTRG.4167.1, where the folding of the was performed using nextPARS experiments as constraints. The color scale indicates the nextPARS score, ranging from red (indicating single-stranded regions) to green (indicating double-stranded regions). **(B)** Three different arc diagrams (R-Chie) are presented. On the left is a comparison of the same MSTRG.4167.1 lncRNA, using nextPARS experiments at 23 degrees (above) and 37 degrees (below). In the middle, the folding with nextPARS experiments at 37 degrees above and 55 degrees below. The last one shows the folding with nextPARS at 23 degrees above and 55 degrees below.

### Comparative analysis of hairpin-like structures reveals enrichment in lncRNAs across Candida species

To further explore the structural landscape of lncRNAs in *Candida* yeast pathogens, we delved into the presence of stem-loop structures essential for the functionality of characterized lncRNAs in diverse species. We noticed that some lncRNAs structures, as determined using nextPARS, presented stem-loop structures. These hairpin-like structures are essential for function in characterized lncRNAs in other species, such as Xist (Maenner et al. 2010), NORAD (Chorostecki, Saus, and Gabaldón 2021; Ziv et al. 2021), LINC00152 (Reon et al. 2018), TCONS_00202587 (Song et al. 2020), roX1 and roX2 (Ilik et al. 2013) among others. We compared the proportion of hairpins in computationally predicted structures of lncRNAs and protein-coding genes in the four *Candida* species (Figure 4A). For this, we scanned for hairpins similar to those previously described on lncRNAs. We observed that the number of hairpin structures normalized by the number of molecules and sequence length was higher in lncRNAs than in protein-coding genes in the four *Candida* species (p-value < 0.0016; Figure 4B). We performed the same approach on humans and *Arabidopsis thaliana* (Figure 4A), and we observed similar results in lncRNAs and protein-coding genes of hairpin structures (Figure 4B). These results suggest that hairpin enrichment in lncRNAs may be a conserved feature of eukaryotes.

**Figure 4.**
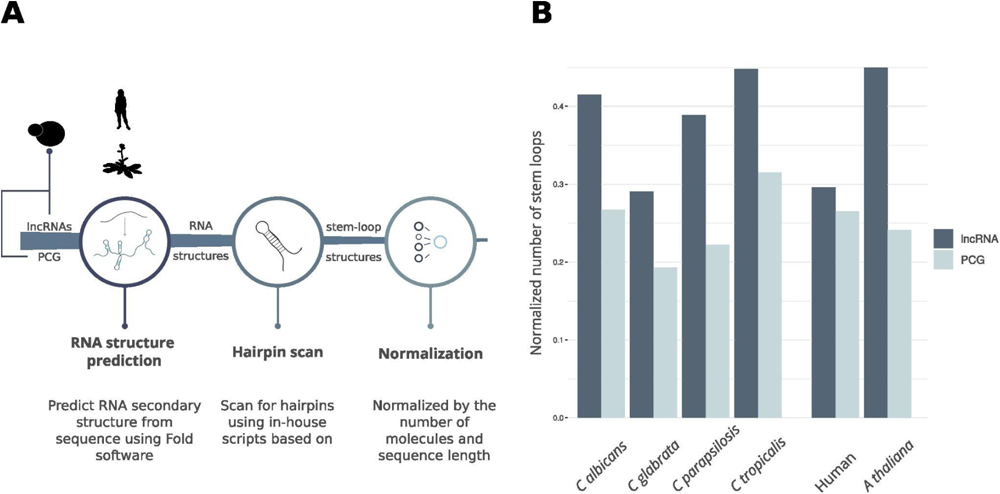
Hairpin comparison between lncRNAs and protein-coding genes. **(A)** Workflow of the pipeline to obtain hairpin structures. **(B)** Barplot showing the comparison between hairpins found in lncRNAs and PCG for the four *Candida* species, *A. Thaliana* and Human

## Discussion

In this study, we performed nextPARS experiments to investigate the structural landscape of mRNAs and lncRNAs in the four major *Candida* pathogens. Our findings provide valuable insights into the relationship between sequence conservation and structural features and the impact of secondary structure on sequence variation. One of the key findings of our study is the correlation between sequence and structure similarity in orthologous mRNAs coding regions. We observed high correlations of nextPARS scores when the sequence identity was above 50%, indicating significant conservation of structural features in highly similar sequences. This correlation was consistent with the phylogenetic relationship of the species, with CTG *Candida* species and post-WGD species belonging to distinct clades. These results highlight the importance of considering both sequence and structure conservation in understanding the functional implications of RNA molecules.

Furthermore, we explored the impact of SNPs on RNA secondary structure. Surprisingly, we found no significant differences between the structural propensities of loci with SNPs and those without SNPs in the coding regions of *Candida* species, contrary to what has been observed for humans (Pegueroles and Gabaldón 2016). However, in the UTRs of *C. glabrata*, SNPs tended to be in less structured positions within the 5’ UTR. This suggests that structural constraints play a role in maintaining the stability and functionality of UTRs, potentially affecting gene expression regulation. These discrepancies in humans and *Candida* lncRNAs may be attributed to species-specific variations in lncRNAs and by the use of high-resolution structural data provided by nextPARS experiments in our analysis, allowing us to uncover subtle structural constraints on sequence variation that might not have been discernible in the context of human lncRNAs. *-*

Comparing the structural characteristics of coding regions and UTRs, we observed that coding regions in *C. glabrata* exhibited significantly higher nextPARS scores than the 5’ and 3’ UTRs. This finding is consistent with previous studies in other species, such as *Saccharomyces cerevisiae* and *Arabidopsis thaliana*. However, it contradicts observations in humans, *Drosophila melanogaster*, and *Caenorhabditis elegans*, where UTRs were reported to be more structured than coding regions. These differences may reflect species-specific variations in RNA structure-function relationships and emphasize the need for further investigations in diverse organisms. It would be worthwhile to explore whether metazoans possess a higher abundance of RNA-binding proteins that interact with CDS, potentially influencing their structural characteristics. Examining this phenomenon using comprehensive annotations could provide valuable insights into the evolution of RNA structures and their regulatory roles in different organisms.

Moreover, our study provides the first experimental characterization of lncRNA structures in the four major *Candida* species. The experimental data revealed the intricate secondary structures of lncRNAs, highlighting their potential independent subdomains and minimal long-range interactions. Using nextPARS data as constraints during RNA folding allowed us to uncover significant differences in certain lncRNAs compared to predictions made without these experimental constraints. Building on this foundational characterization, we focused on the specific lncRNA, MSTRG.4167.1, revealing temperature-dependent folding variations. The observed differences in structural conformations at different temperatures shed light on this particular lncRNA. These findings contribute to our understanding of *Candida* lncRNAs and underscore the importance of incorporating experimental data to enhance the accuracy of *in-silico* predictions in studying RNA structural landscapes. Interestingly, we identified hairpin-like structures in lncRNAs, which are functionally important in other species. Comparing the proportion of hairpin structures between lncRNAs and protein-coding genes, we found that lncRNAs were enriched in hairpin structures in *Candida* species, as well as in humans and *Arabidopsis thaliana*. One possible explanation for this phenomenon could be that these hairpin structures in lncRNAs contribute to their stability and function as structural scaffolds, aiding in interactions with other molecules or proteins within the cell. These structural features might play crucial roles in processes such as RNA-protein interactions, localization, or regulation of gene expression, highlighting the importance of further investigation into the functional implications of these hairpin-like structures in lncRNAs.

Our study comprehensively analyzes the structural landscape of *Candida* mRNAs and lncRNAs, revealing important insights into the relationship between sequence, structure, and function. The observed correlations between sequence and structure conservation and the differences in structure between coding regions and UTRs highlight the intricate interplay between RNA sequence and structure in gene expression regulation. The observed enrichment in hairpin structures of lncRNAs suggests their potential functional significance in diverse organisms. Further investigations into RNA structure and evolution will deepen our understanding of the intricate gene expression mechanisms and provide possible therapeutic strategies against *Candida* infections.

## Author Contributions

TG and UC conceived and designed the study; ES performed the experiments; UC analyzed the data and developed the algorithms; UC and TG wrote the manuscript. All authors read and approved the final manuscript.

