## Supplementary material for "Genome-wide RNA structural determination in *Candida* yeast pathogens": Supplementary material.pdf

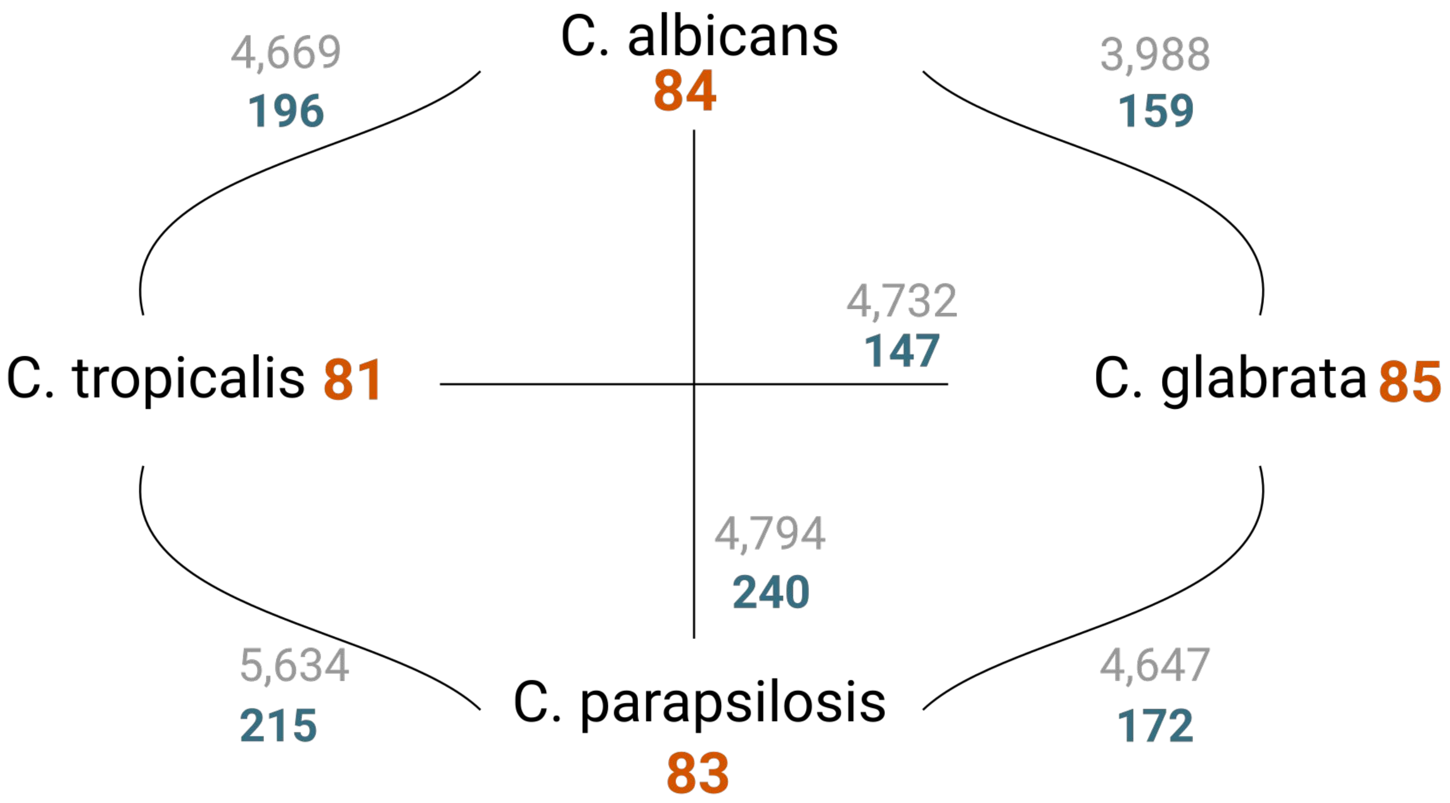


**Figure S1.** Genes with orthology information in the 4 *Candida* species

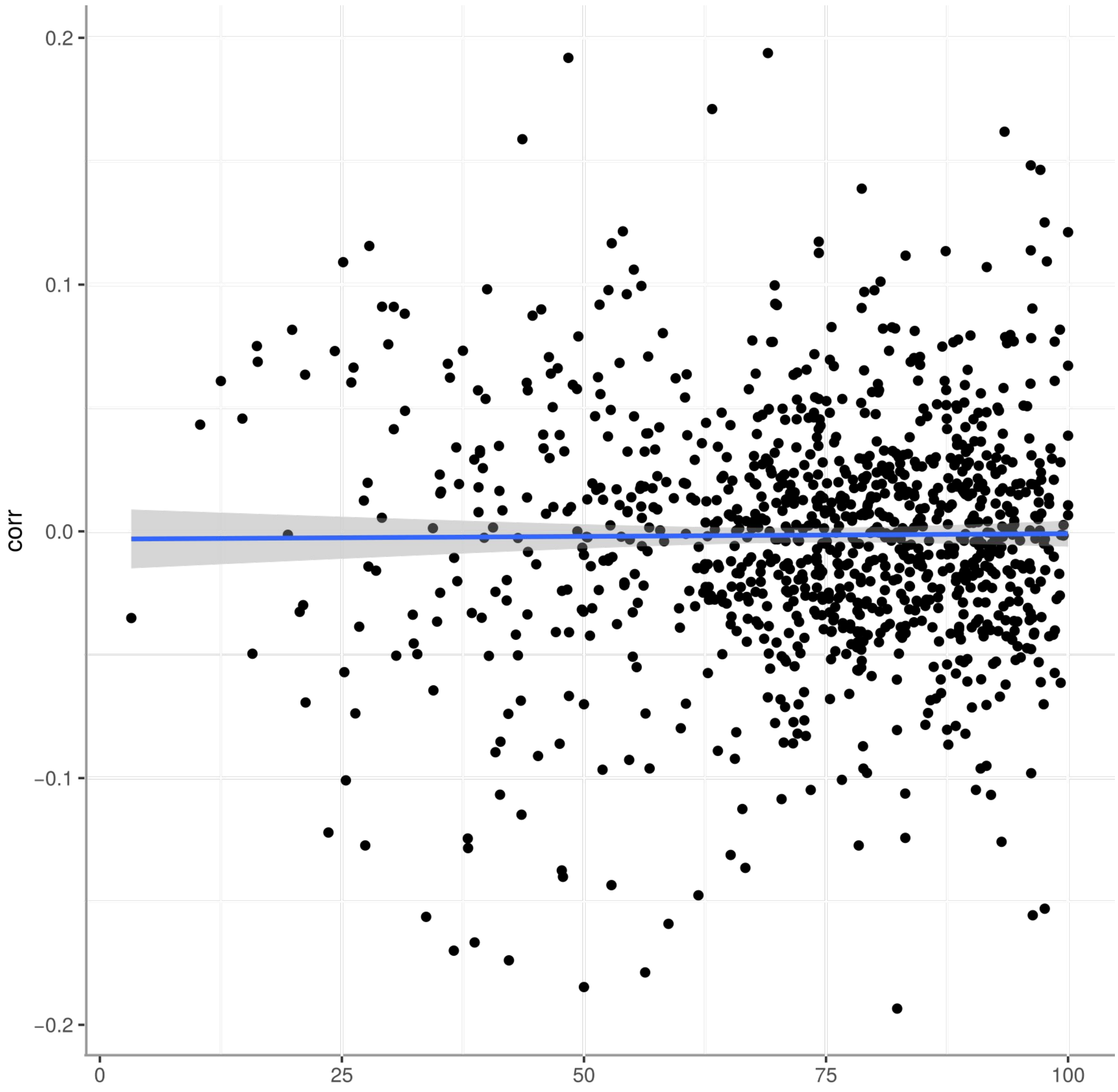


**Figure S2.** Correlations between randomly shuffled scores

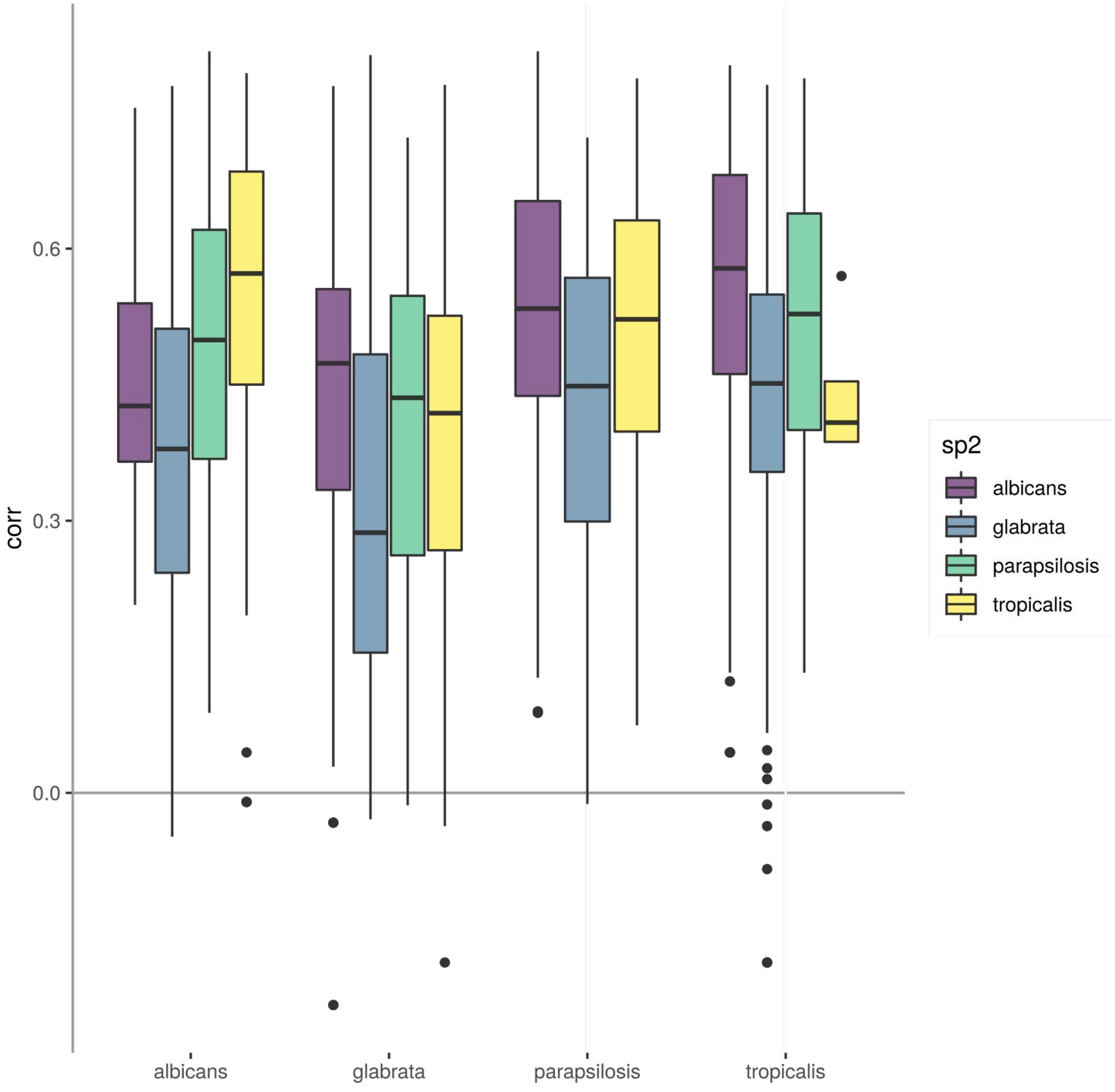


**Figure S3.** Correlation between sequence and structure in orthologs for nextPARS score in aligned positions

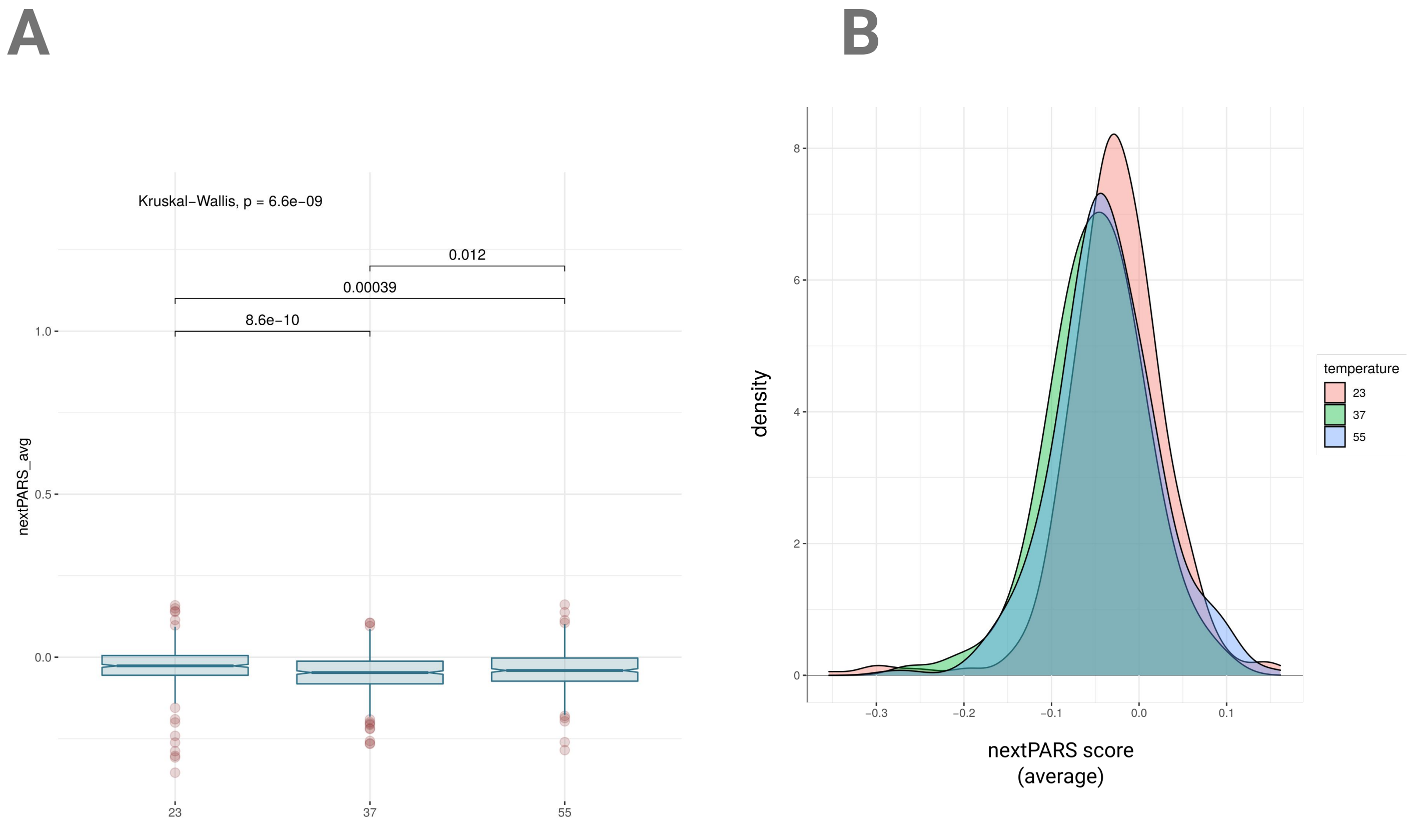


**Figure S4.** C Parapsilosis at different temperatures. **(A)** Box plot of nextPARS score for C. parapsilosis mRNAs at different temperatures. **(B)** Density plot of nextPARS score comparing conserved positions against non-conserved ones at three different temperatures.

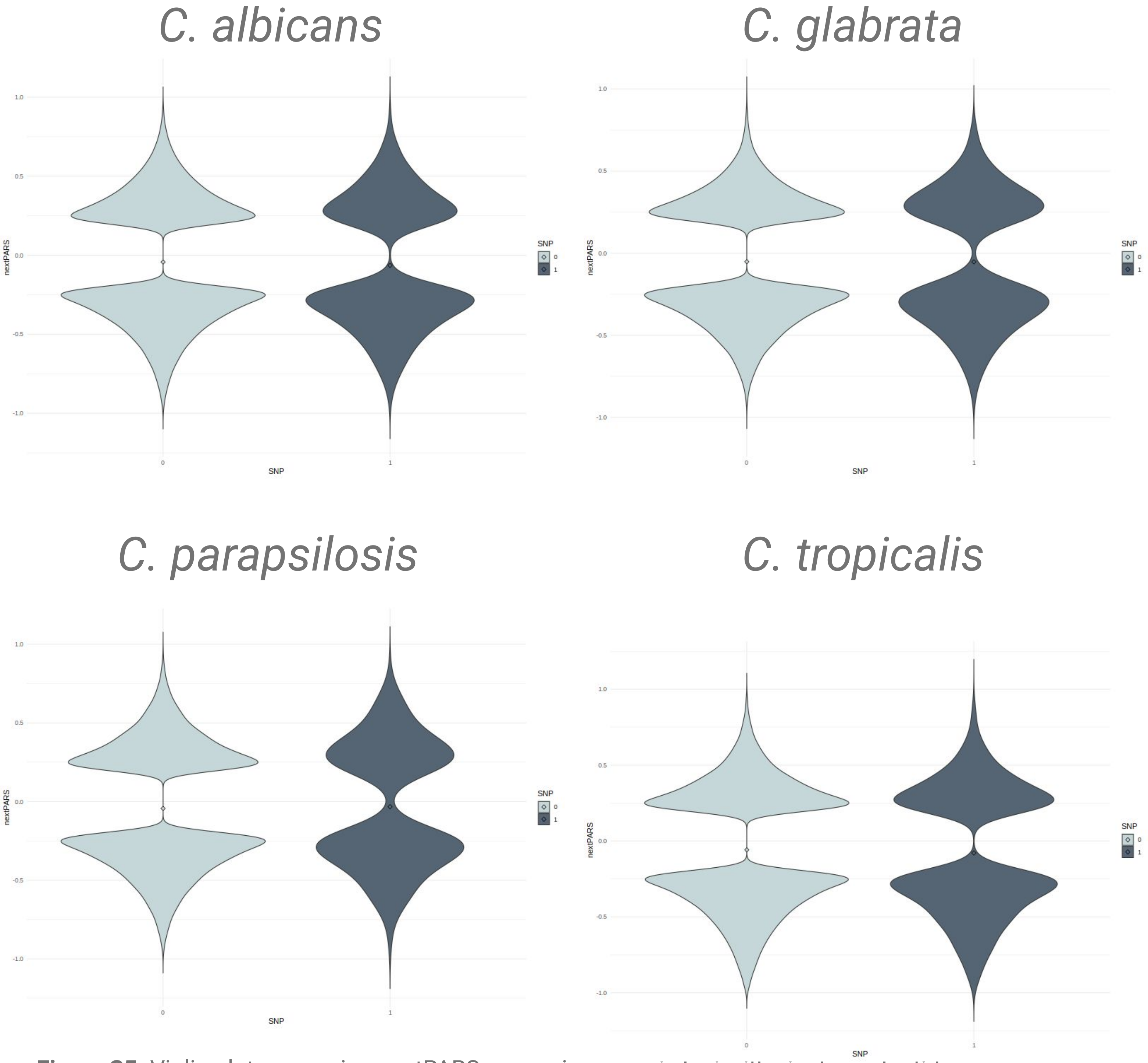


**Figure S5.** Violin plot comparing nextPARS scores in genomic loci with single nucleotide polymorphisms (SNPs) to those without reported SNPs across four *Candida* species.

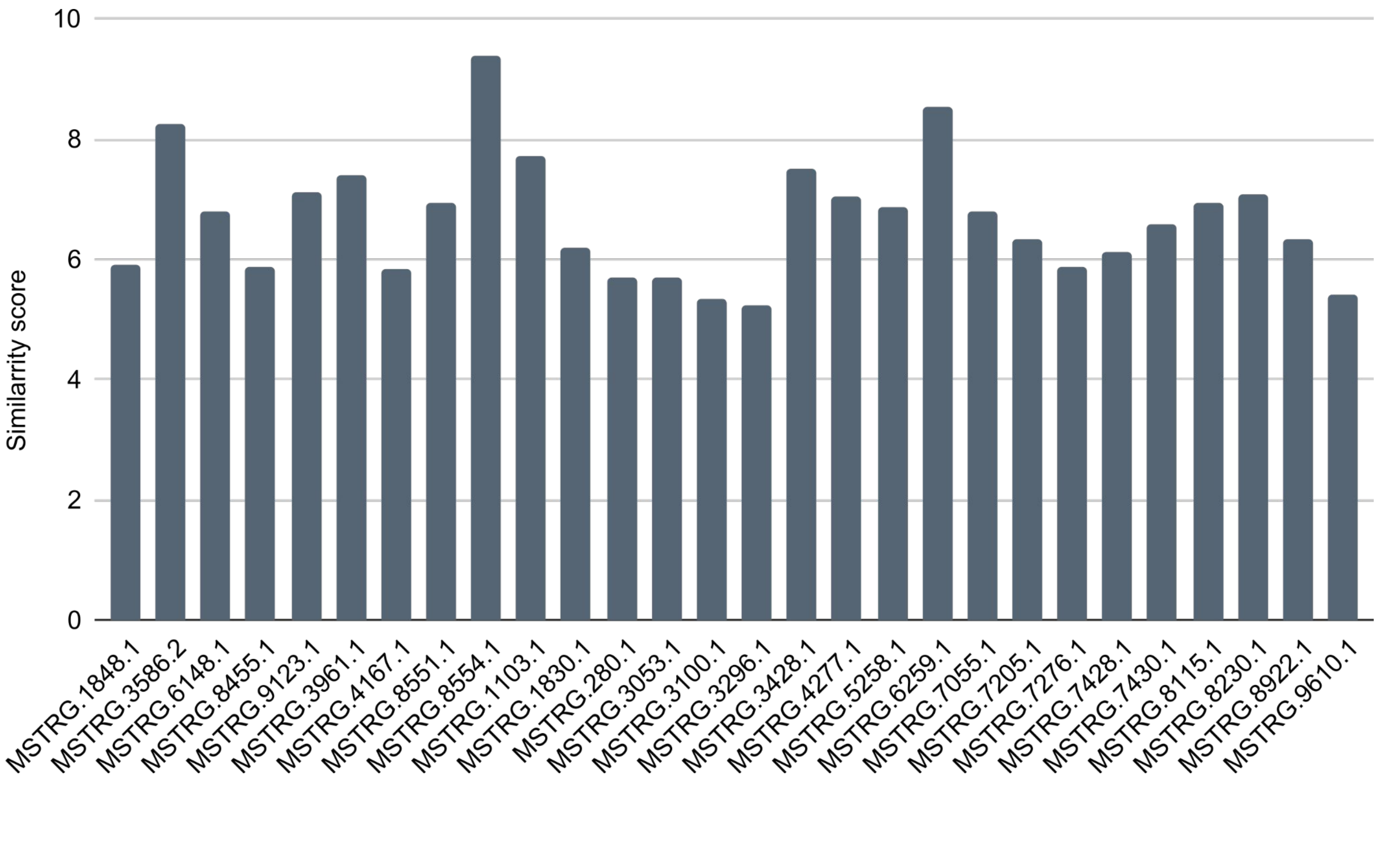


**Figure S6.** Bar plot illustrating the similarity scores for various long non-coding RNAs (lncRNAs), comparing the scores obtained with experimental constraints (nextPARS data) to predictions made without these constraints.

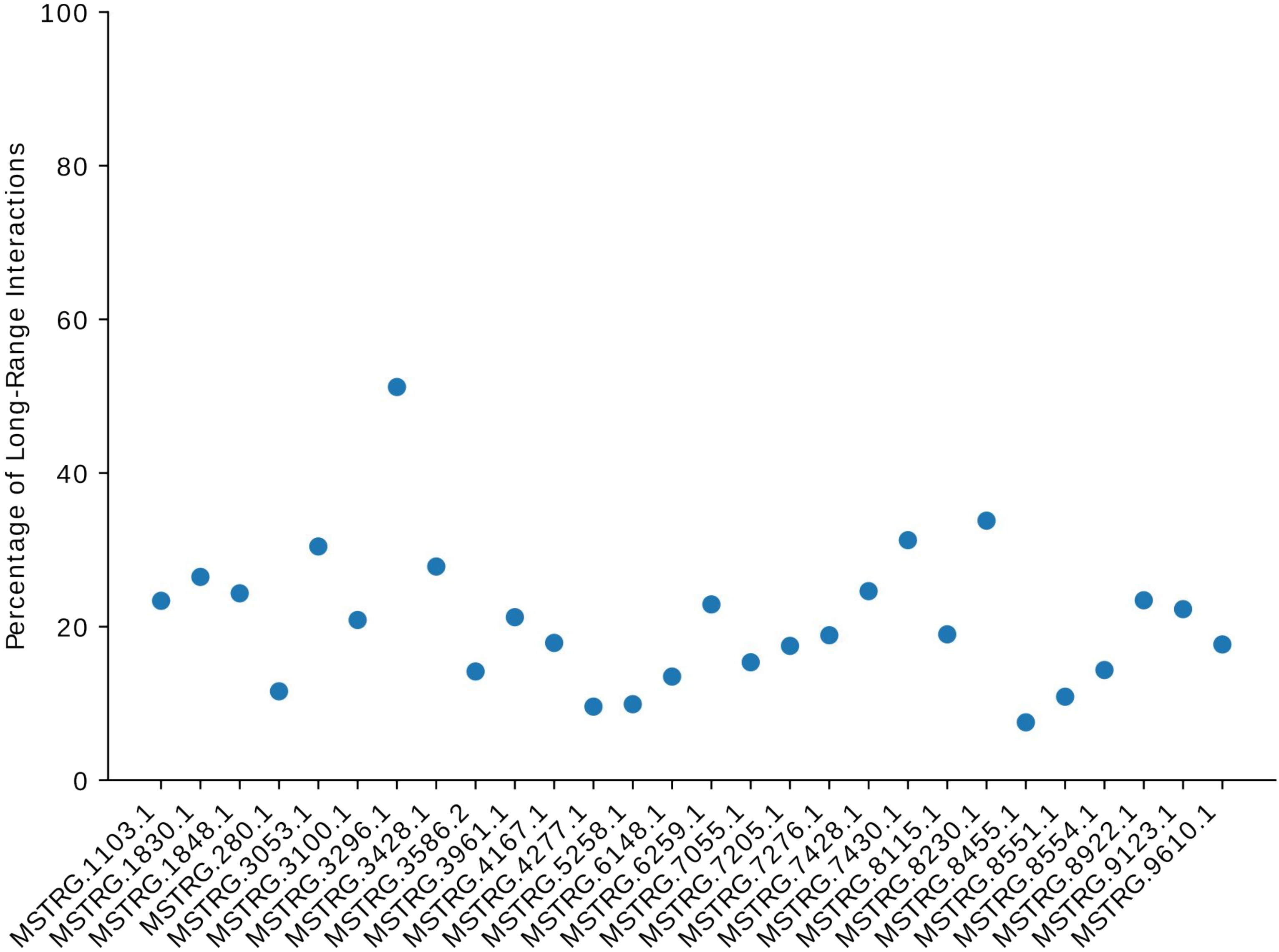


**Figure S7.** Visual representation of the distribution of long-range interactions across the analyzed lncRNA sequences. The x-axis of the plot corresponds to the unique identifier of each lncRNA, while the y-axis represents the calculated percentage of long-range interactions. The resulting plot provides

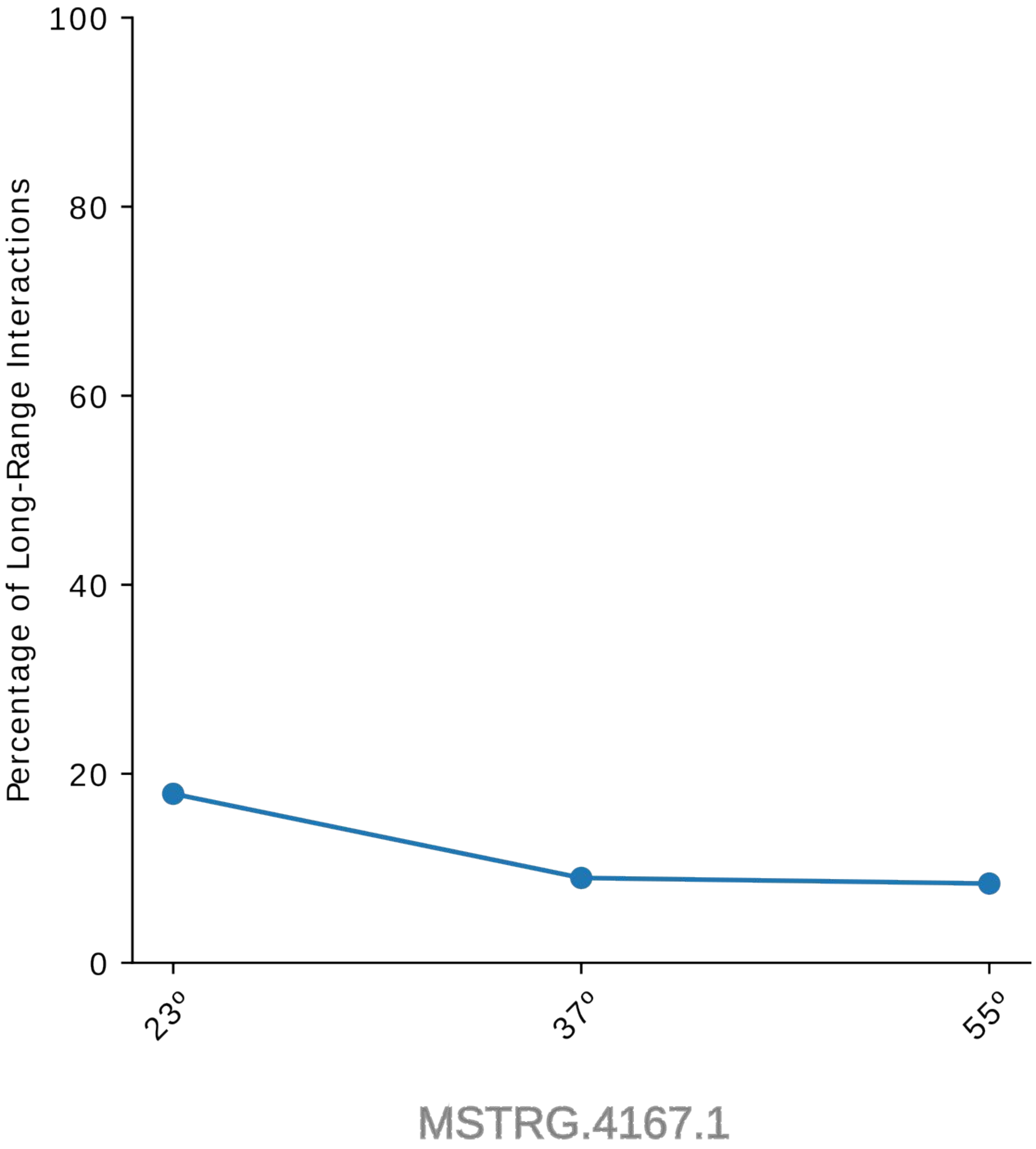


**Figure S8.** Percentage of long-range interactions of MSTRG.4167.1 lncRNAs at three different temperatures

| | *C. albicans* | *C. glabrata* | *C. parapsilosis* | *C. tropicalis* |
|---|---|---|---|---|
| **PCG** | 286 | 296 | 412 | 285 |
| **lncRNAs** | 5 | 1 | 3 | 19 |

**Table S1.** Number of mRNAs and lncRNAs detected with sufficient confidence (> five average counts per position) in the four *Candida* species

| MSTRG.4167.1 | 23º vs 37º | 37º vs º55 | 23º vs 55º |
|---|---|---|---|
| **Similarity score** | 6.634375 | 6.164139 | 5.88634 |

**Table S2.** Similarity scores for the structural comparison of lncRNA MSTRG.4167.1 at different temperatures (23°C, 37°C, and 55°C).
